# Sexual selection for males with beneficial mutations

**DOI:** 10.1101/2021.05.25.445661

**Authors:** Gilbert Roberts, Marion Petrie

## Abstract

Sexual selection is the process by which traits providing a mating advantage are favoured. Theoretical treatments of the evolution of sex by sexual selection propose that it operates by reducing the load of deleterious mutations. Here, we postulate instead that sexual selection primarily acts through females preferentially mating with males carrying beneficial mutations. We used simulation and analytical modelling to investigate the evolutionary dynamics of beneficial mutations in the presence of sexual selection. We found that female choice for males with beneficial mutations had a much greater impact on genetic quality than choice for males with low mutational load. We also relaxed the typical assumption of a fixed mutation rate. For deleterious mutations, mutation rate should always be minimized, but when rare beneficial mutations can occur, female choice for males with those rare beneficial mutations could overcome a decline in average fitness and allow an increase in mutation rate. We propose that sexual selection for beneficial mutations could overcome the ‘two-fold cost of sex’ much more readily than choice for males with low mutational load and may therefore be a more powerful explanation for the prevalence of sexual reproduction than the existing theory. If sexual selection results in higher fitness at higher mutation rates and if the variability produced by mutation itself promotes sexual selection, then a feedback loop between these two factors could have had a decisive role in driving adaptation.

## 1. Introduction

Mutation is the source of variation and the overwhelming majority of mutations are deleterious (i.e they have a negative impact on fitness) [1]. Because of this, most models have ignored beneficial mutations (i.e. those with a positive impact on fitness), deeming them too rare to be of interest (see [2] for an exception). Nevertheless, adaptation depends upon those rare occasions when mutations have a beneficial impact on fitness, especially in changing environments [3, 4]. Here, we investigate the role of sexual selection in favouring beneficial mutations. Sexual selection can be a powerful process resulting in strongly biased mating success [5]. This can even allow modifiers of the mutation rate (‘mutator genes’, such as factors controlling DNA repair [6, 7]) to persist, because female choice selects for those males with beneficial mutations [8, 9]. We postulate here that female choice can be so potent that it not only promotes the maintenance of genetic variation, but it allows beneficial mutations, despite their comparative rarity, to have a marked impact on fitness. This means that the two-fold cost of sex may be overcome not just by the lower reduction in fitness caused by deleterious mutation load [10, 11], but by the increase in fitness resulting from fixation of new beneficial mutations. Our focus in this paper is therefore on how sexual selection has a more important effect than has previously been considered in favouring beneficial mutations and driving adaptation, as opposed to the widely considered effect of decreasing deleterious mutational load. We further postulate that female choice can result in higher fitness at higher mutation rates, despite the decline expected from the predominance of deleterious mutations, because female choice is effective in selecting those males with an increased number of beneficial mutations. Any explanation for the evolution of sexual reproduction within groups must show how the production of males, even when they do not care for offspring, increases the genetic quality of sexually produced offspring, to overcome the numerical reduction of reproducing offspring, the ‘two-fold cost of sex’ [12, 13]. Sexual reproduction can entail variance in mating success, especially where females can choose between males of varying quality and where a male can mate with more than one female [14, 15]. This differential mating success is integral to the process of sexual selection and it has previously been suggested that this process could contribute to the maintenance of sex by the selective removal of low quality males from the breeding population [10, 11]. Sexual selection could thereby reduce the risk of population extinction [2], as has been demonstrated in flour beetles *Tribolium castaneum* [16]. According to this theory, females can pick those males with the lowest load of deleterious mutations and those males that do not contribute to the breeding population are effectively a sink for deleterious mutations.

We simulated the ‘genetic quality’ of individuals by examining the evolution of deleterious and beneficial mutations in asexual and in sexual populations, and by varying the degree of female choice in the latter. Implicit in our model is the assumption that, in addition to determining survival, the mutations that a male accumulates determine the condition of some trait, which is used by female subjects in mate choice. Thus, we assume that a male’s genetic quality is revealed in the trait and the female subjects use that information to select the best male. Choice thereby functions to identify the highest quality mates [3, 17]. This is supported by the literature on ‘good genes’ effects in sexual selection [18, 19] and by the demonstration that sexual traits can reveal genetic quality [20-22]. It is also consistent with other sexual selection models [23]. We compared sexual and asexual populations in the presence of varying levels of sexual selection and a modifier of mutation rate. We hypothesized that increasing mutation rate above a baseline would leverage the effects of sexual selection making it more likely for sexual types to have higher genetic quality than asexual types. The rationale for this is that an increase in mutation rate feeds variation in genetic quality (and hence attractiveness), and that this variation promotes choice between males [15]. We consider that sexually reproducing individuals will vary in mating success [11] and that a key driver of this variation is female choice for high quality and attractive mates that will provide females with high viability and attractive offspring. We propose that females can actually get ‘good genes’ rather than ‘fewer bad genes’ as in the models of how sexual selection facilitates the evolution of sex [10, 11]. As such, we predicted that variability in mutation rate should facilitate the effect of sexual selection in overcoming the two-fold cost of sex.

## 2. Methods

We simulated the ‘genetic quality’ of individuals by examining the evolution of deleterious and beneficial mutations in asexual and in sexual populations, and by varying the degree of female choice in the latter. A simplified flow diagram of the simulations is given in Figure S6 and a Visual C program is provided as supplementary information. Our simulation methods were based on models of the evolution of mutation rate [6, 8]. As in those simulations, the parameters reflected literature estimates where available [24], subject to the constraint that the model was intended as a simple abstraction of reality and not an attempt to simulate an entire genome.

The simulations began by setting up a population of *P* individuals. In simulations of sexual populations, exactly half were male and half female. Individuals were given a pair of homologous ‘chromosomes’ (i.e., we assumed diploidy) each bearing a ‘mutator gene’ and an associated string of 10 ‘viability genes’. We did not assume that these genes constituted the entire genome; only that there were no interactions between the genes of interest and those at other sites and that for the purposes of comparison between simulations, all other things were equal.

Each viability gene was subjected to a mutation process which could introduce deleterious and/or beneficial mutations. By default, deleterious mutations occurred with probability 10^−3^ per gene per generation and beneficial mutations at 10^−6^ per gene per generation, so deleterious mutation occurred at a rate 1000x that of beneficial mutations. This implemented both the principle that there are many more ways to introduce faults in a complex organism than there are to improve it; and the empirical finding that beneficial mutations are much rarer than deleterious ones. We note that estimates of the per nucleotide mutation rate for sexually reproducing species tend to be around 1 × 10^−8^ per individual per generation [24-26]). However, if we were to use literature mutation rates we would also need to use realistic numbers of genes because selection acts at the level of the individual carrying those genes. This would then effectively mean we were trying to simulate entire genomes. We cannot do this, nor is this what the model is for. Instead, what we try to do is provide a ‘proof of concept’ by modelling a scenario where individuals are subject to selection based on differences in numbers of mutations. We assumed that each mutation had a sexually concordant effect. A mutator gene increased the rates of beneficial and of deleterious mutation by a factor *M*. We assumed that the mutator gene affected DNA repair only in a relatively small region of the genome [27], namely the set of viability genes referred to above; that it was adjacent to the first of the row of viability genes; and that the crossover rate between the mutator and the first viability gene was the same as between each other viability gene.

Individuals were subjected to a mortality process, whereby their probability of survival was a function of their genetic quality. Deleterious mutations were assumed to have larger phenotypic effects than beneficial mutations: deleterious mutations reduced the wild type fitness of 1 by 0.005; beneficial mutations increased it by 0.002. The model assumed co-dominance with additive fitness effects, so the effects of mutations were summed to give ‘genetic qualities’, which determined individual survivorship in a mortality process.

Surviving individuals reproduced, either by asexual or by sexual reproduction, the latter with or without female choice. Reproduction replaced the population by selecting females at random as parents, each such selection producing one offspring, with or without a male, which was either selected at random or chosen from a set of *n*. Asexual reproduction was implemented by selecting an individual at random and copying its chromosomes into an individual in the next generation. Sexual reproduction without female choice involved selecting a male and a female subject at random. In the case of female choice, a female subject was selected at random, and a set of *F* male subjects was selected at random. The female then bred with the male of highest genetic quality from that set of *F* males. This ‘best of n’ rule is the most widely used rule in modelling female choice [28, 29] and has some empirical support from lekking species [30]. For each sexual mating, one offspring was produced by bringing together chromosomes contributed by both parents. This was carried out by selecting a chromosome at random from each parent and allowing crossover between each adjacent gene with probability 0.01. The process of selecting parents and producing offspring was repeated until the population was replaced by a new generation of *P* individuals.

Each scenario was simulated 10 times, allowing for uncertainty estimates to be computed across runs which each produced different results due to the stochastic processes in the models. Unless otherwise stated, parameters used were as given in Table S1.

## 3. Results

Considering first the results with the default mutation rates, if we compare Figure 1 a and b, we can see that female choice had a marked effect in decreasing deleterious mutations (means and standard errors across 10 simulations at generation 100 were 1.7526+-0.4349 and 0.04200 +- 0.06530 with no female choice and with female choice between two males respectively) and in increasing beneficial mutations (from 0.0003+-0.0055 to 0.2000 +- 0.1897). We can approximate the relative effects of female choice on beneficial mutations as increasing them by a factor of 0.2000/0.0003 = 667, as compared to decreasing deleterious mutations by a factor of 1.7526/0.042 = 42. Figure 1 shows that this difference continued to increase markedly with the number of generations.

**Figure 1.**
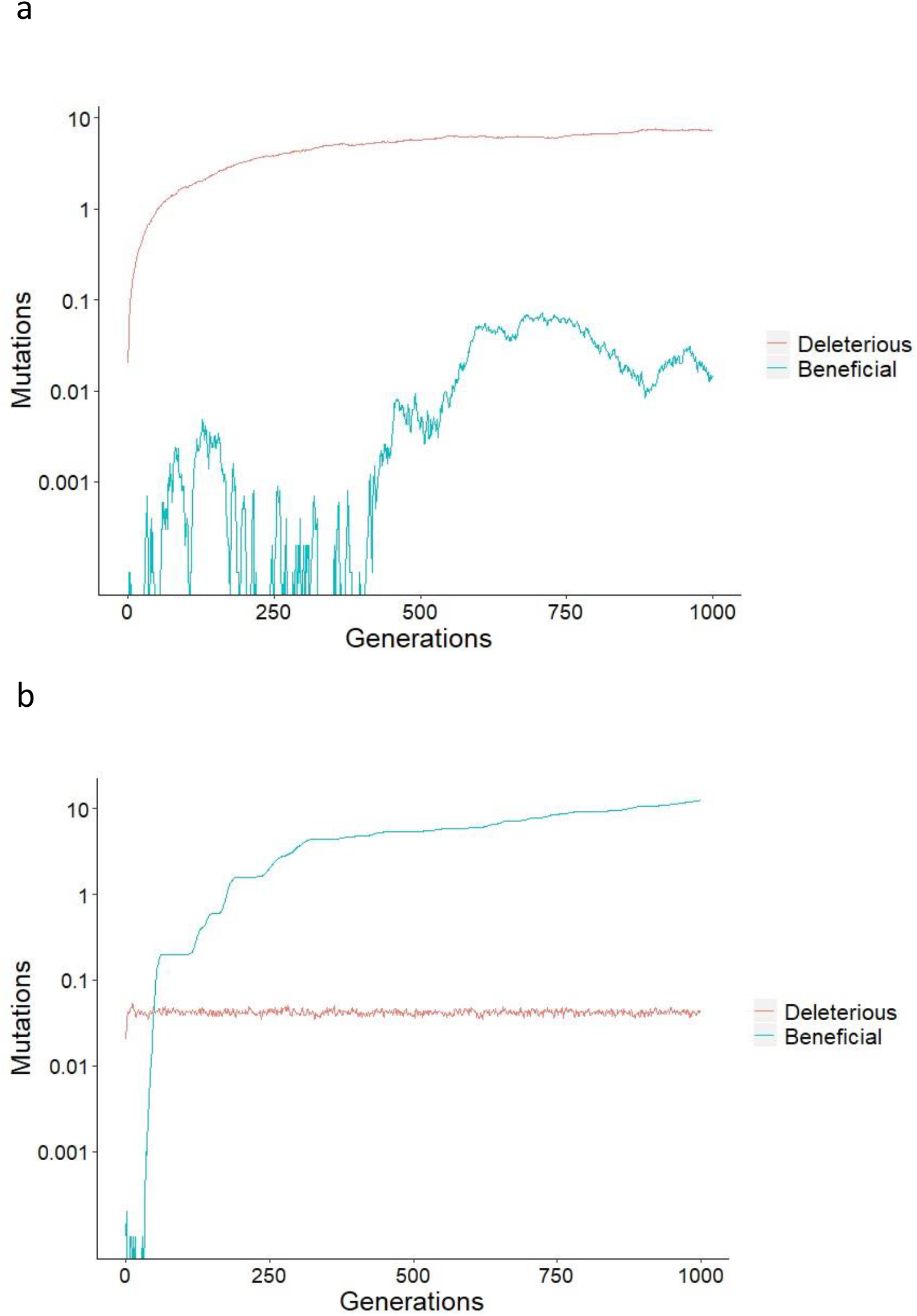
Simulated evolutionary dynamics of the numbers of deleterious and beneficial mutations with (a) no female choice and (b) female choice between two males. Plotted are the means across 10 simulations. Parameters were: population size N=1000, equally divided between males and females; Rate of deleterious and of beneficial mutations per gene per generation = 10^−3^ and 10^−6^ respectively; effect of deleterious and beneficial mutations on default fitness of 1 = 0.005 and 0.002 respectively; cost of female choice = 0.02*female choice.

Looking at the relationship between mutation rate and deleterious mutations, we found that in populations with asexual reproduction and in those with sexual reproduction but no sexual selection, deleterious mutation load increased dramatically when mutation rate increased (Figure 2a). However, female choice, even between just two males, reduced deleterious mutations to very low levels. Female choice was so effective that even with a high mutation rate, the numbers of deleterious mutations were reduced to substantially below those found in asexual populations or in sexual populations lacking female choice. Conversely, numbers of beneficial mutations were low, even with a ten-fold increase in mutation rate in asexual populations and sexual ones without female choice (Figure 2b). However, with female choice, beneficial mutations were much more common, increasing from 12.49 ± 1.52 through 15.47 ± 1.57 to 17.19 ± 1.48 with female choice of 1, 2 and 5 males respectively; and an increase to ten times the mutation rate accentuated this effect (with 3.29x, 5.87x and 6.38x the numbers of beneficial mutations with female choice of 1, 2 and 5 males respectively). Summing these effects, overall genetic quality decreased with increasing mutation rate in asexual populations and sexual ones lacking female choice, yet it increased with increasing mutation rate in populations with female choice (Figure 2c). Thus, sexual selection was so powerful that it overcame the 1000-fold disadvantage (see Methods) of the beneficial mutation rate and allowed an increase in genetic quality with increased mutation rate. This increase in genetic quality occurred because sexual selection was effective at keeping numbers of deleterious mutations low despite an increase in mutation rate, yet was also effective in selecting for beneficial mutations, and could do this most effectively when mutation rate was high.

**Figure 2.**
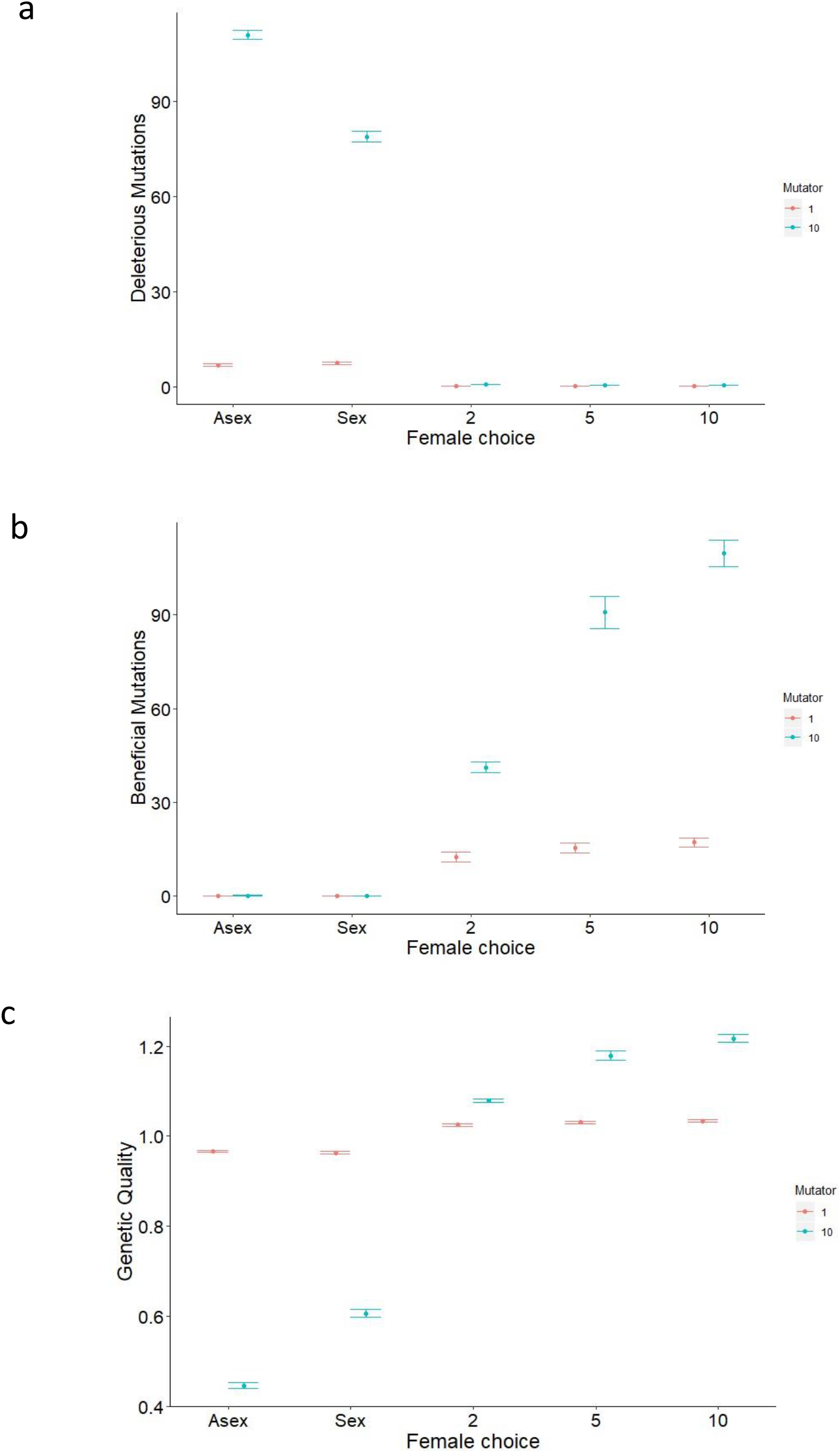
The figure shows how the numbers of mutations per individual vary with mode of reproduction and strength of female choice where “Asex” is asexual reproduction; “Sex” is sexual reproduction without female choice; and where “2”, “5” and “10” are sexual reproduction with female choice between the given number of males. Figures a and b show the numbers of mutations per individual; c shows ‘genetic quality’ calculated as baseline fitness (1) minus the number of deleterious mutations per individual multiplied by the effect of each (0.005); plus the number of beneficial mutations multiplied by the effect of each (0.002). The rate of deleterious mutations per gene per individual per generation was 1000x that of beneficial mutations. Plotted are the means and standard errors computed across 10 simulations for each set of parameters, taken at generation 1000. Plotted in red are the results where the mutator gene *M* = 1; blue is where *M* = 10, i.e. the rates of both deleterious and beneficial mutations are at 10x the baseline mutation rates, which were 10^−3^ and 10^−6^ respectively. Other parameters: population size N=1000, equally divided between males and females in simulations with sexual reproduction; cost of female choice = 0.02*female choice.

We further investigated the interaction between female choice and mutation rate (Figure 3). Genetic quality increased with mutation rate and female choice to an intermediate maximum before declining rapidly. High levels of female choice improved genetic quality; but there came a point where even the highest level of female choice shown was insufficiently powerful not to be overwhelmed by the surge in deleterious mutations caused by a substantially increased mutation rate.

**Figure 3.**
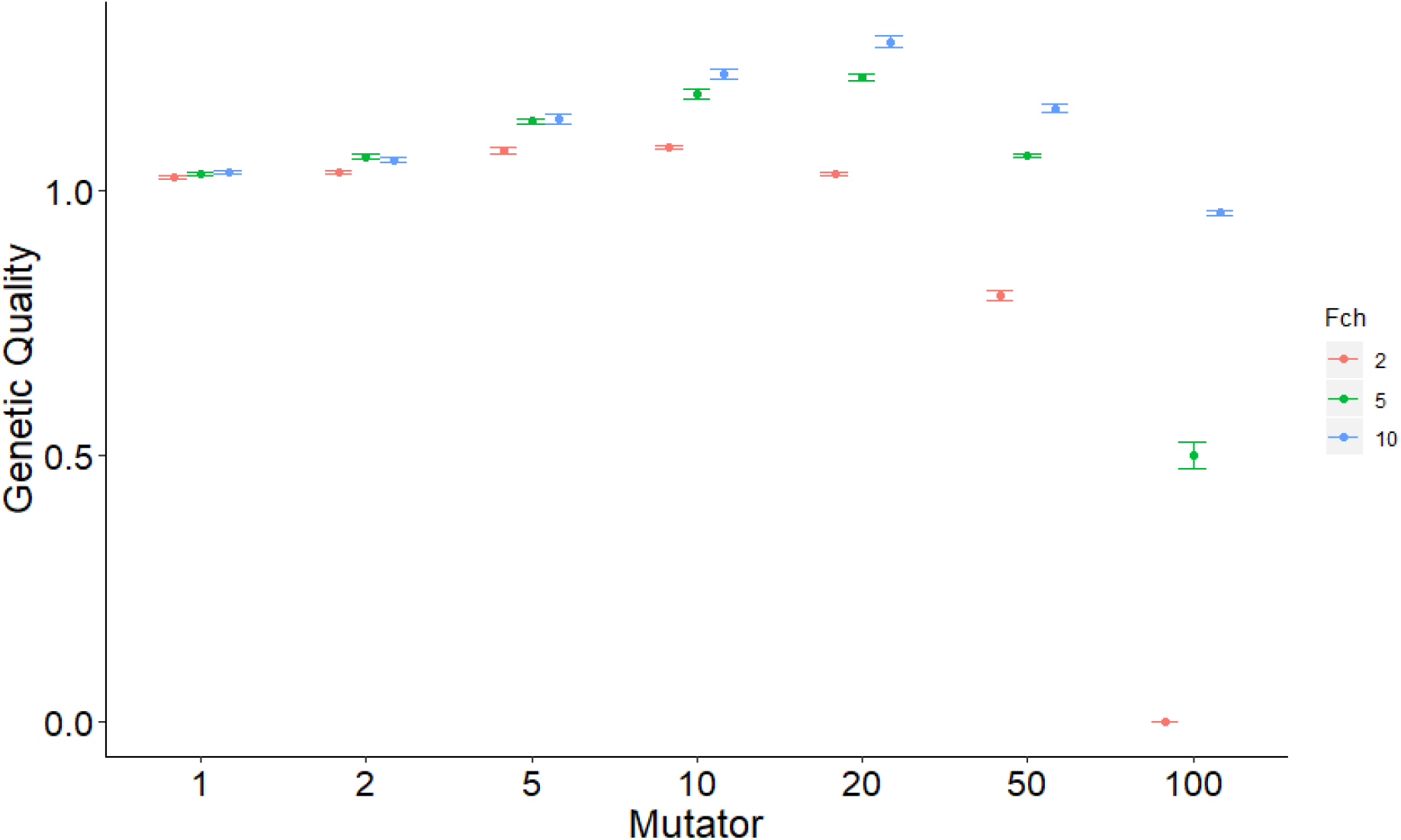
The figure shows how ‘genetic quality’ varies with a ‘Mutator’ and with sexual selection by female choice. Genetic quality was calculated as given in Fig. 2. The mutator was a multiplier of the default mutation rates. Female choice was between 2, 5 and 10 males. Plotted are the means and standard errors computed across 10 simulations for each set of parameters, taken at generation 1000. Other parameters were as in Fig. 2.

We investigated the robustness of our results by varying key parameters whilst maintaining a moderate degree of female choice (*F* = 5). First, we varied the ratio of deleterious to beneficial mutation around the default ratio of 1000:1 respectively. As intuitively expected, genetic quality increased as beneficial mutations became relatively more common, and especially when this factor combined with a higher mutation rate (Figure S1). Looking at the effect of each beneficial mutation on genetic quality, we can again see that the results are intuitive (Figure S2): increasing the effect of beneficial mutations increases genetic quality, especially when mutation rate is high. Varying the cost of female choice within the range shown has no effect on genetic quality in either mutation rate condition (Figure S3). Increasing the population size, *P*, increased the effect size, as expected from the increased potential for genetic change across more individuals (Figure S4). Varying recombination rate had a little quantitative effect, with increased recombination allowing an increase in genetic quality, presumably through allowing favourable genetic combinations (Figure S5). All in all, varying key parameters suggested that the main effect we describe was robust.

The benefit of simulations is that they can help us predict processes that are difficult to comprehend intuitively or mathematically; the corollary is that they can be hard to interpret. To better understand the processes in the model we hypothesized that female choice selects for a breeding population that results in offspring that are *s* standard deviations above the mean of the underlying population. To test whether males that were chosen to breed were indeed of higher genetic quality than the population from which they were drawn, we aggregated across the first 1000 generations for all 1000 reproducing pairs and calculated the mean difference in genetic quality between females and their chosen males. Where female choice was absent and mutation was at the default rate, the mean difference was 0.0001 ± <0.00005 (standard error of the mean); with females choosing the best of 10 males and mutation rate at 10x the default, the difference was 0.0022 ± <0.00005. Therefore, females were choosing males that had a genetic quality equivalent to approximately 1 beneficial mutation above the average (where each beneficial mutation had an effect of 0.002 on fitness, as represented by the 10 simulated genes). In this way, sexual selection appears to be able to exploit increasing variation caused by increased mutation rate and actually produce an increase in genetic quality out of a background that one would otherwise expect to be dominated by increased deleterious mutation load.

As a simple analytical approximation, consider that female choice between males results in offspring that are *s* standard deviations above the mean genetic quality. For these offspring to be of a quality that overcomes the two-fold cost of sex we need:

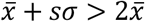

And therefore:

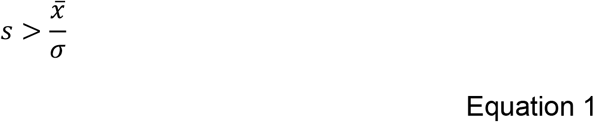

This equation will most readily be satisfied when the standard deviation of genetic quality is high relative to the mean. That is, there needs to be high variability within the population. Variability results from mutation, and we can show when an increase in mutation rate can be favoured. This can occur when the decrease in the mean genetic quality in the population that necessarily results from an increase in mutation rate (assuming there is a strong predominance of deleterious over beneficial mutations) is more than compensated for by the effect of female choice in selecting for males of high genetic quality. If subscripts 1 and 2 indicate the means and standard deviations before and after the change in mutation rate, then:

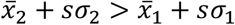

So:

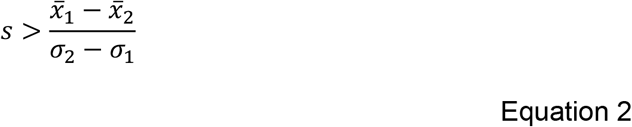

This analytical approach has the benefit that it considers the whole genome, something we do not attempt in our simulations, which allow us to include stochastic factors at the level of the gene. As a hypothetical numerical example, consider that increasing the mutation rate lowers the mean genetic quality from 1 to 0.9 whilst increasing the variance in genetic quality from 0.2 to 0.4. Consider also that female choice selects for males that produce offspring 1 standard deviation above the mean. Entering these hypothetical values into Equation 2 demonstrates that an increase in mutation rate combined with female choice can indeed lead to an increase in the genetic quality of offspring. To aid understanding, this example is illustrated in Figure S6. Whilst simplified, this makes the point that an increase in mutation rate can be favoured under sexual selection and can thereby contribute to overcoming the two-fold cost of sex.

## 4. Discussion

We have shown here that sexual selection is a potent force not just in reducing deleterious mutation load but in leveraging the spread of beneficial mutations. Several empirical studies have found that sexual selection over relatively short periods of time (typically under 100 generations) resulted in increased average fitness or reduced extinction [31]. Intuitively, it seems more plausible that these results are observed because sexual selection is acting on the already present standing variation, which largely comprises deleterious alleles. However, our result shows that sexual selection for beneficial mutations can actually be the more important effect, even over moderate numbers of generations and even when beneficial mutations were 1000 times rarer than deleterious ones and when they each had only 40% of the impact on fitness. We suggest that rather than males being effectively a sink for bad genes [10, 11] they can be a vehicle for good genes.

The power of sexual selection was also apparent when we allowed for an increase in mutation rate. We found that sexual selection for beneficial mutations could result in an increase in genetic quality with increased mutation rate. This is counterintuitive given that any increase in mutation rate brings 1000 times as many deleterious as beneficial mutations in our model. An increase in mutation rate can never be adaptive in a model that considers only deleterious mutations; it is only by allowing for rare beneficial mutations that we can discern this effect. The result highlights just how powerful the effect of female choice can be, even when just choosing between 2 males. Not only can sexual selection combined with increased mutation rate provide a solution to the paradox of how variation can be maintained [8, 32], but it can also lead to an increase in genetic quality.

We believe this is the first time that this effect has been reported. Its significance is that if mutation rate is elevated above a typically postulated minimum level then this would increase the benefits of sex relative to asexual reproduction, helping to overcome what has been termed the ‘two-fold cost of sex’ [12, 13]. It therefore seems that the role of sexual selection in the promotion of heritable genetic variation is key to understanding the predominance of sexual reproduction.

Our model employs ‘best of n’ female choice as a convenient way of modelling sexual selection. However, this is just one way of implementing variance in mating success in which some individuals take a disproportionate share of mating opportunities. Active choice by females is clearly something that must have evolved after sexual reproduction itself, but in line with theoretical treatments that have focussed on deleterious rather than beneficial mutations [10, 11], we posit that some variance in mating success is inevitable and that this could have contributed to the invasion of sexual reproduction in an asexual population in the first instance. Current thinking suggests that anisogamy has evolved from isogamy [33, 34]. The evolution of anisogamy needs to be accompanied by an effect on mating success. Only when an increase in mating success of smaller male gametes overcomes the cost of loss of a male resource contribution to the zygote will this system be evolutionarily stable. Where mating success is determined by the level of indirect genetic benefits, the male phenotype will evolve to reveal their underlying genetic quality and only when the genetic quality differences are large and constantly maintained by mutation will this system be evolutionarily stable [8, 15, 35]. Signalling of genetic quality by males must therefore have evolved either concurrently or very early on in the evolution of anisogamy.

Our model assumes that mutation can be controlled genetically. One mechanism for this is through selection on those genes that are responsible for DNA repair [6], but there are other possible genetic mechanisms [36]. We predict a greater mutation rate in genes that influence sexually selected phenotypes and in more sexually selected species. Across non-human species, particularly in birds, which differ in within population variance in mating success [37], and thus the level of sexual selection, there is some evidence of a positive correlation with the rate of mutation, measured by variance in minisatellite mutation rate [38, 39]. Given that sexual selection can often result in greater variance in mating success in males, our results are also consistent with male-biased mutation rates [40]. Our results may also be consistent with the findings of experimental work on seed beetles [41]: interestingly, an experiment reported that not only did sexually selected males pass on a lower mutation load but that they had fewer *de novo* mutations suggesting that that sexual selection interacts with mutation rates.

We have shown here that sexual selection can cause fitness to be higher when mutation rate is higher; but we have not shown that this can dynamically drive an increase in mutation rate through individual level selection. To do this would require an understanding of how much individuals gain themselves from having a high mutation rate and how much they gain from others having a high mutation rate. These fitness benefits depend in turn on with whom individuals interact, which means considering population size and structure. Hence to limit the complexity of the analysis presented we have here confined ourselves to considering fitness benefits at the population level from fixed mutation rates.

Our results suggest a positive feedback effect whereby a high mutation rate could favour sexual reproduction over asexual reproduction, while sexual reproduction with sexual selection could favour a high mutation rate. We speculate that this could create a ratchet effect in that once sexual reproduction is established, it would become harder to switch back to asexual reproduction. We predict that the transition from sexual reproduction to asexuality should be particularly rare in populations with strong sexual selection and that parthenogenetic females could only invade a population of sexually reproducing females when there is little or no genetic variation in male genetic quality. This could occur in small, isolated inbred populations that become genetically depauperate. There is some evidence that parthenogenetic populations can occur on islands [42]. Interestingly, some island populations of birds are also known to lose the sexually selected signals that are characteristic of mainland populations [43].

Natural selection will tend to favour a low mutation rate [1], yet adaptation requires mutation. Our model shows how a faster rate of adaptation can occur with sexual selection. This is consistent with the finding that sexual selection (measured as the degree of polygyny) interacts with the rate of molecular evolution and with body mass to predict species richness at the genus level [44]. We suggest that evolvability itself is under selection [45], and that through sexual reproduction with sexual selection, evolution can lead to greater evolvability.

## Author contributions

GR devised the computer simulations and analysis and wrote the first draft of the paper. MP conceived the original idea of relating sexual selection and mutation rate and contributed to the theoretical background.

