## Supplementary Table and Figures for "Sexual selection for males with beneficial mutations"

**Sexual selection for rare beneficial mutations promotes the evolution of sexual reproduction and adaptation**

**Supplementary Information**

**Table S1. Parameters, abbreviations and their defaults.**

| **Parameter** | **Default** |
| --- | --- |
| Population size (*N*) | 1000 |
| Number of males and females in sexual populations | 500 |
| Generations | 1000 |
| ‘Mutator’ (*M*) – multiplicative modifier of mutation rate | 1 |
| Female choice (*F*) | 1 |
| Cost of female choice | 0.02 x *F* |
| Ratio of deleterious to beneficial mutations | 1000 |
| Rate of deleterious mutations per gene per generation | 1 x 10^-3^ |
| Fitness | 100 |
| Effect of deleterious mutations | -0.5 |
| Effect of beneficial mutations | 0.2 |
| Rate of recombination between genes | 0.01 |

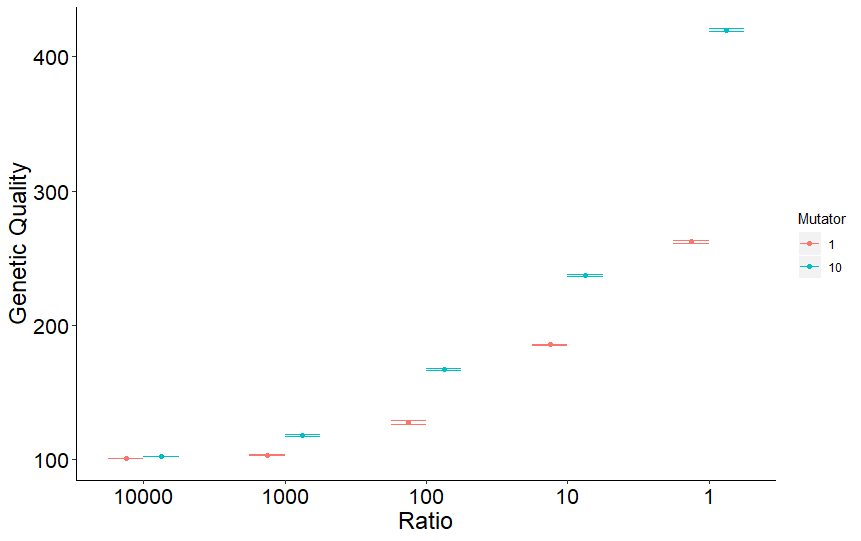

**Figure S1.**

Genetic quality (as in Fig. 1) in relation to the ratio of deleterious to beneficial mutations (varied around the default of 1000) with the Mutator set to 1 or 10. Other parameters as in Fig. 1.

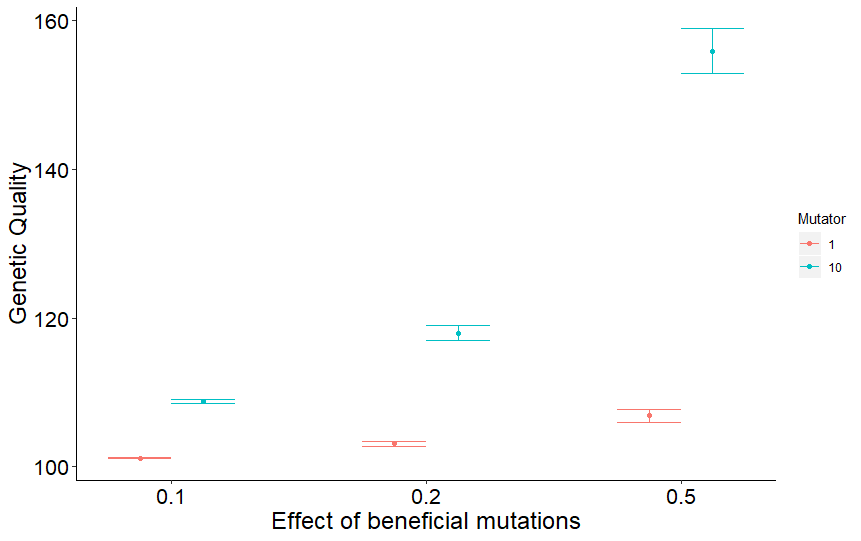

**Figure S2.**

Genetic quality (as in Fig. 1) in relation to the effect of beneficial mutations (varied around the default of 0.2) with the Mutator set to 1 or 10. Other parameters as in Fig. 1.

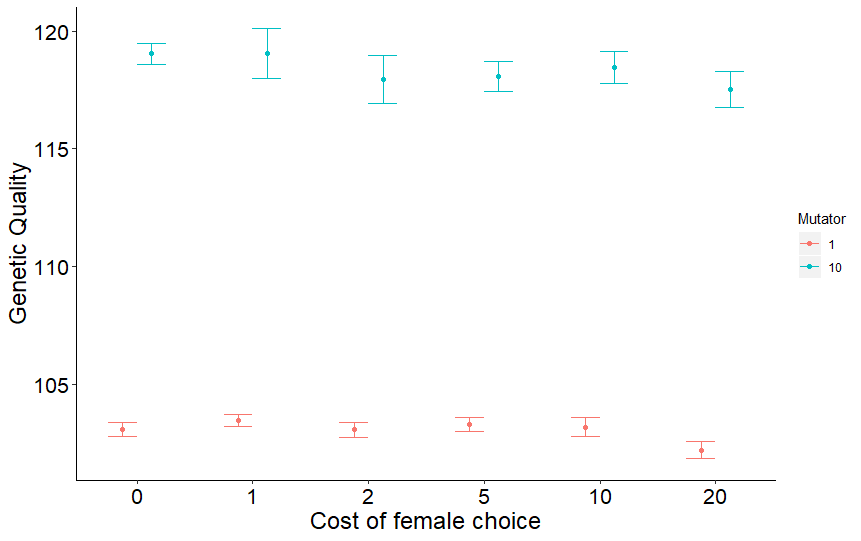

**Figure S3.**

Genetic quality (as in Fig. 1) in relation to the cost of female choice (varied around the default of 2 x the value of the female choice parameter *F*) with the Mutator set to 1 or 10. Other parameters as in Fig. 1.

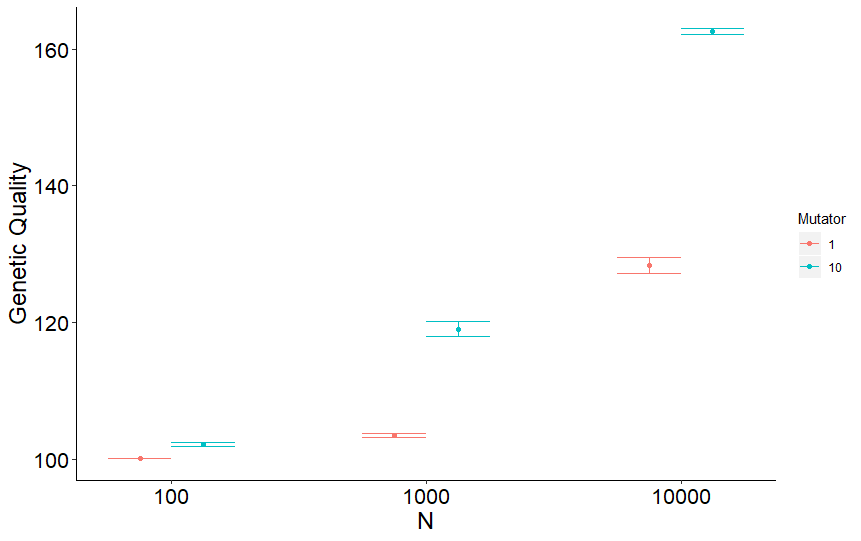

**Figure S4.**

Genetic quality (as in Fig. 1) in relation to population size, *N*, varied around the default of 1000 and with the Mutator set to 1 or 10. Other parameters as in Fig. 1.

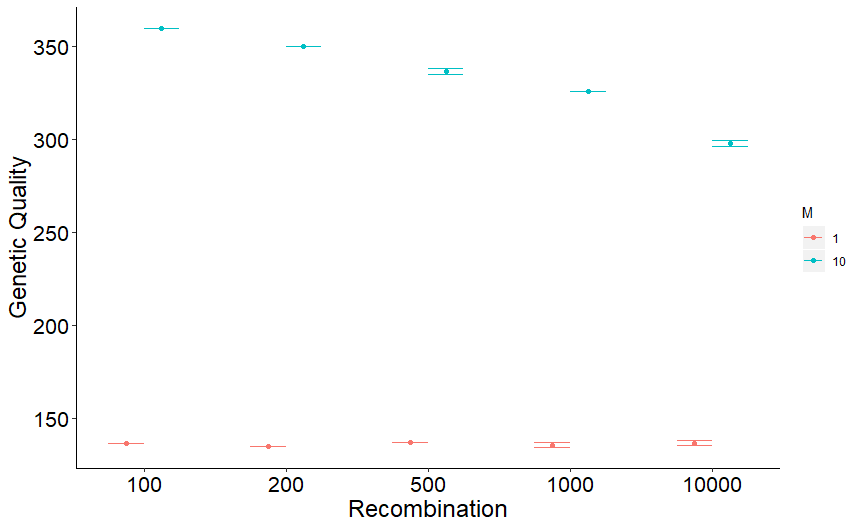

**Figure S5.**

Genetic quality (as in Fig. 1) in relation to recombination rate (displayed as the reciprocal), showing the effect of decreasing recombination from the default of 0.01, with the Mutator set to 1 or 10. Other parameters as in Fig. 1.

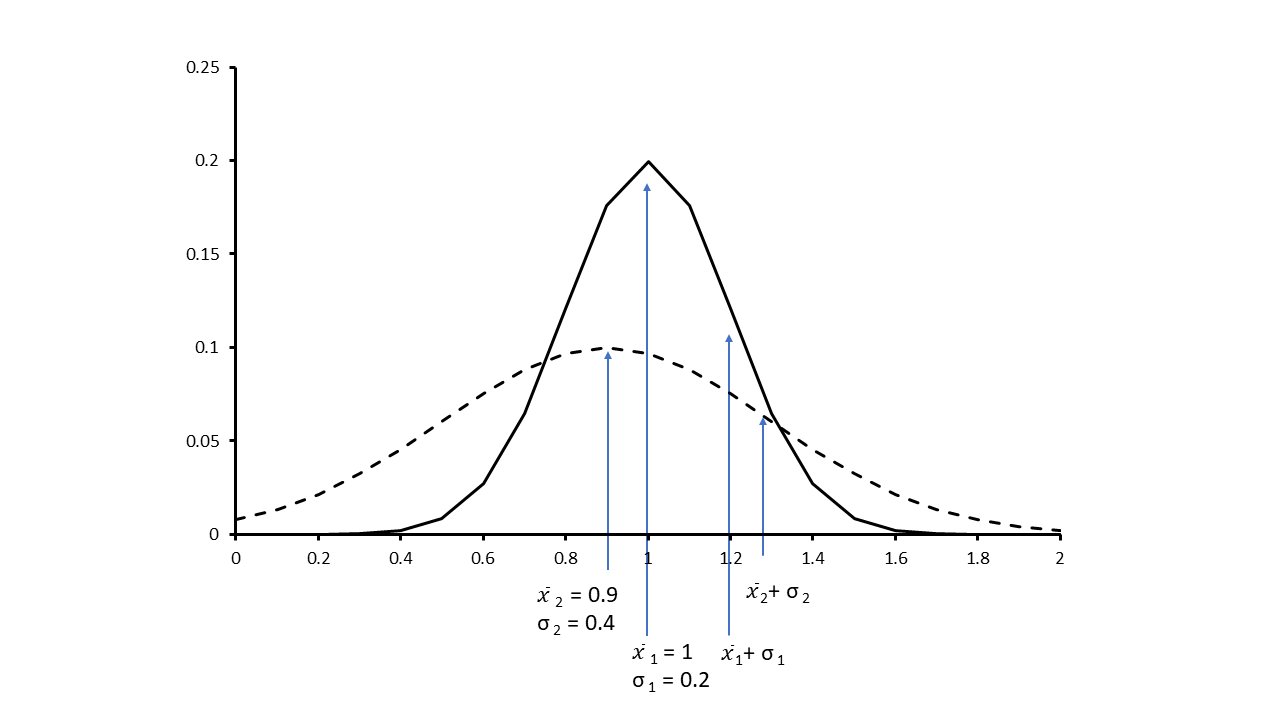

**Figure S6.** To illustrate the hypothetical case where a decrease in mean is accompanied by an increase in variance, meaning that the sample mean one standard deviation above the new population mean can actually increase.

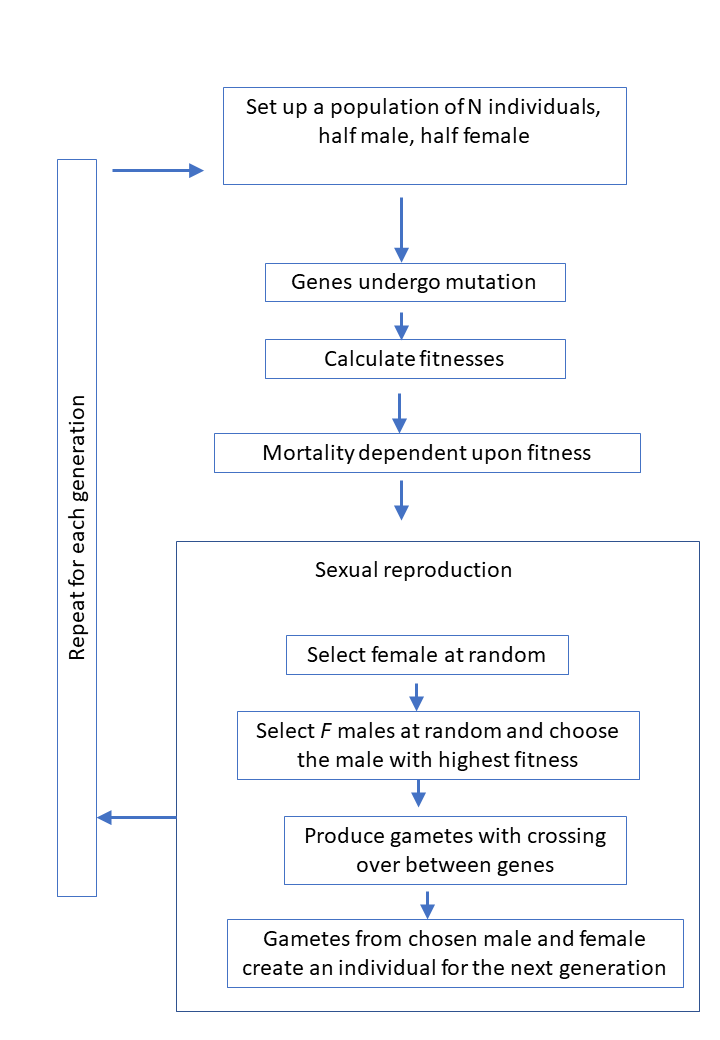

**Figure S7.** Flow diagram showing a simplified representation of the simulation program for sexual populations with female choice.
